# Reciprocal exchange obscures phylogenetic patterns of introgression

**DOI:** 10.1101/2025.09.10.675447

**Authors:** Devin Y Mendoza, Evan S Forsythe

## Abstract

Hybridization and introgression are major evolutionary forces in eukaryotes that have shaped the evolutionary trajectories of crops, livestock, and modern humans. Yet current theoretical and statistical frameworks provide only a partial view of the dynamics and genomic consequences of introgression. An unexplored dimension is the possibility that introgression between two lineages can occur simultaneously in both directions at a single genomic locus—a process we term *reciprocal introgression*. Although this type of introgression is plausible, especially in cases of bidirectional introgression across the genome, it remains largely undetectable under widely used statistical approaches, hindering evaluation of its prevalence in natural systems. Moreover, unrecognized reciprocal introgression may bias summary statistics commonly used to quantify gene flow, obscuring our ability to resolve introgression events. To address this, here, we develop theoretical expectations of the impact of reciprocal introgression on phylogenetic and site-pattern-based results. To test these expectations, we implement coalescent simulations to model genomes experiencing reciprocal introgression and apply sliding-window introgression statistics to these data. We find that reciprocally introgressed loci produce phylogenetic patterns opposite to those expected under unidirectional introgression. These mixed signals can dilute evidence of introgression or even generate false signals of introgression events that never occurred. Taken together, our results highlight the need for updated statistical approaches and pave the way toward a more complete understanding of the impacts of introgression on gene histories and organismal evolution.

## Introduction

Hybridization is widespread across eukaryotes (Mallet 2005; Yakimowski and Rieseberg 2014; Mallet et al. 2016; Taylor and Larson 2019). Hybridization, followed by backcrossing, leads to introgression of alleles between divergent species or populations, a phenomenon which has been well-documented in plants (Rieseberg and Soltis 1991; Rieseberg 2006; Suarez-Gonzalez et al. 2016; Forsythe et al. 2020a) and animals (Dasmahapatra et al. 2012), including humans (Green et al. 2010; Prüfer et al. 2014; Kuhlwilm et al. 2016). The growing availability of full genome sequences has led to the development of statistical approaches for detecting patterns of historical introgression (Green et al. 2010; Durand et al. 2011; Martin et al. 2015; Pease and Hahn 2015; Hahn and Hibbins 2019; Hibbins and Hahn 2021). The D-statistic or “ABBA-BABA test” is a site-pattern-based test that was developed to detect an abundance of derived alleles shared by non-sister species, which serves as evidence that the two species underwent introgression (Green et al. 2010; Durand et al. 2011). While this concept has been expanded upon (Martin et al. 2015; Martin and Jiggins 2017; Zheng and Janke 2018; Dagilis et al. 2021), the D-statistic remains a widely used tool in identifying and quantifying introgression in genomic datasets.

An important aspect of introgression that is not resolved with the D-statistic is the direction of introgression, which is critical for understanding the impact of introgression because novel traits are acquired by the recipient lineage but not the donor. Several statistical methods have been developed to determine the direction of introgression (Pease and Hahn 2015; Hibbins and Hahn 2019, 2021; Forsythe et al. 2020b). However, these statistics largely assume that all introgressed loci across the genome were introgressed in the same direction (*i*.*e*., unidirectional introgression) and provide a summary statistic that indicates the prevailing direction of introgression across the genome. A shortcoming of these methods is that they are unable to resolve cases in which loci across the genome were introgressed in opposite directions (*i*.*e*., bidirectional introgression). Bidirectional introgression has been demonstrated to occur in nature (Yaakub et al. 2006; Wang et al. 2016; Pazmiño et al. 2019), but its detection has largely relied on population genetic, demographic, and biogeography information, meaning that studies focused primarily on site-pattern and phylogenetic data may underestimate its prevalence.

Given the general absence of tools to explore bidirectional introgression, the potential effects of introgression directionality on biasing D-statistics and related methods are underexplored. While it is established that a locus introgressed in one direction will exhibit a similar D-statistic to a separate locus introgressed in the opposite direction (Durand et al. 2011), little consideration has been given to a scenario in which a single locus experiences introgression in both directions, thus resulting in mutual displacement. Here, we explore this possible novel mode of introgression and use coalescent simulations to define the expected effects on phylogenetic and site-pattern-based analyses, including its potential to bias and obscure genomic studies of introgression.

## Results

### Phylogenetic expectations under alternative introgression scenarios

We used pipe diagrams to outline the expected gene tree topologies of a locus undergoing alternative introgression scenarios (Fig. 1). It is well understood that introgression in either the P2->P3 or P3->P2 direction produces gene trees and site patterns uniting P2 and P3 (Fig 1A and B). This phylogenetic expectation assumes that an incoming allele displaces the native allele at a locus. However, it is also conceivable that incoming alleles could displace native alleles in both directions, a scenario which we describe as ‘reciprocal introgression’. Our pipe diagram predicts that this type of introgression will yield a gene tree topology in which P1 and P3 are sister and share the derived allele (Fig. 1C). Under the typical four-taxon introgression framework, this so-called “BABA” site pattern would only be expected under introgression between P1 and P3 or incomplete lineage sorting; however, our model suggests a novel scenario that could conceivably yield such gene tree topology/site patterns even when introgression only occurs between P2 and P3.

**Figure 1:**
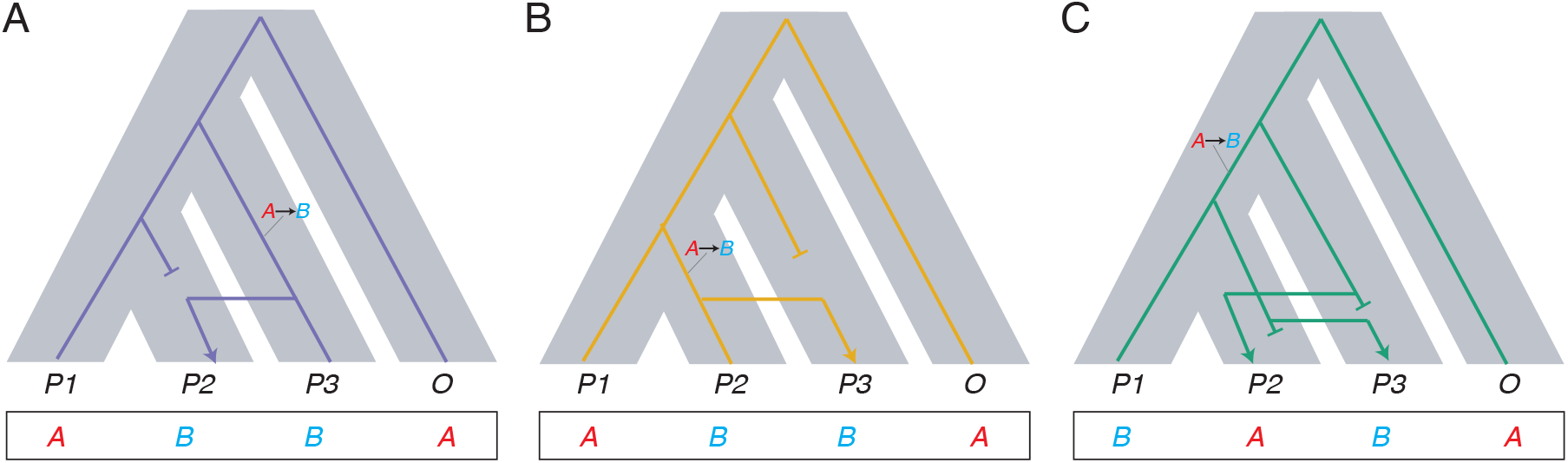
Expected gene tree topologies and site patterns under alternative introgression scenarios. Pipe diagrams showing the background species topology (thick gray branches) and the topology of gene trees that underwent introgression from P3 to P2 (A; purple branches), P2 to P3 (B; yellow branches), or reciprocal introgression (C; green branches). Blunt end branches indicate alleles displaced by an introgressed allele (arrow). Phylogenetically informative nucleotide substitutions (“A”: ancestral state; “B”: derived state) are shown along branches and the resulting site patterns are shown at the bottom.

### Simulations of reciprocal introgression

We simulated a chromosome alignment with tracts of introgression in both directions between P2 and P3, with overlapping genomic regions amounting to reciprocally introgressed tracts (Fig 2A). We calculated the D-statistic for 1,000 non-overlapping windows (Fig. 2A, bottom) and plotted genome-wide distributions (Fig. 2B). As expected, D-statistics calculated from non-introgressed tracts are centered on zero and unidirectional introgression tracts exhibit positive D-statistics. Importantly, as predicted in Figure 1C, reciprocal introgression tracts exhibit a clear skew toward negative D-statistics (Fig. 2). Despite the clear negative shift in the reciprocal introgression distribution, it is worth noting that some windows exhibited near-zero D-statistics. We observed this unexpected pattern to varying degrees across 100 replicate runs (Table S1), likely representing the stochastic nature of coalescent simulations (Discussion). Taken together, our results highlight a novel mode of introgression between P2 and P3 that mimics the genomic signature of introgression between P1 and P3.

**Figure 2:**
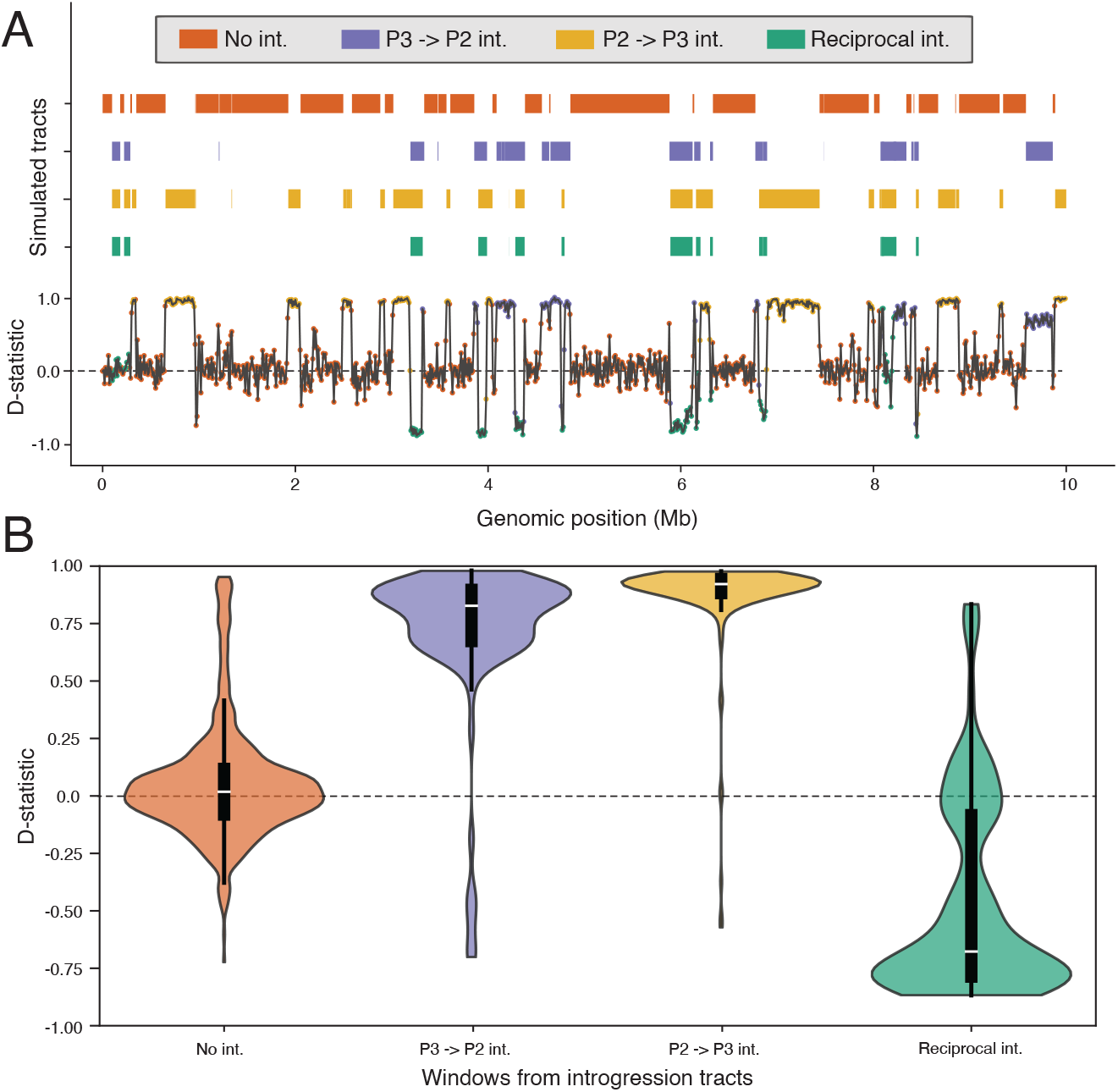
The genomic landscape of D-statistics under reciprocal introgression. (A) Simulated chromosome with tracts (i.e. haplotype blocks) that underwent alternative introgression histories. D-statistic estimates are shown for windows spanning the genome, which colored points indicated the introgression history of the window. (B) Violin plots showing the distributions of D-statistic estimates across windows of alternative introgression types. Box plots show medians and quartiles from distributions.

## Discussion

### Interpreting mixed distributions in introgression landscapes

Our theoretical work and simulations define reciprocal introgression as a novel scenario in which introgression between a single pair of lineages can produce both positive and negative D-statistics. These findings introduce two major implications for interpreting four-taxon introgression results. First, genome-wide counts of ABBA and BABA site patterns are often used to calculate a single D-statistic estimate that summarizes the introgression pattern across the genome. This type of genome-wide summation is used to assess the significance and extent of introgression based on the relative frequencies of ABBA and BABA sites. However, given that reciprocal introgression introduces an excess of BABA sites (Fig 2), this behavior would act to dilute the overall excess of ABBA across the genome, leading to underestimates of introgression in genome-wide D-statistic values. For example, the simulated genome in Figure 2 yielded a genome-wide D-statistic of D=0.22. However, removing reciprocally introgressed windows from the estimate yields a D-statistic of D=0.29, demonstrating that reciprocal introgression dilutes the magnitude of D-statistic estimates.

Secondly, sliding window D-statistic and related F-statistic (Martin et al. 2015) analyses are frequently used to identify the genomic loci that were introgressed between species. In some cases, these analyses revealed the surprising result that some genomic blocks exhibit positive D-statistics, while neighboring blocks exhibit negative D-statistic (Crawford et al. 2015; Zhang et al. 2016). This type of D-statistic sign mosaicism has been interpreted as evidence that multiple gene flow events have occurred between different pairs of taxa. However, our analyses demonstrate that reciprocal introgression can mimic these patterns. Given this result, it is imperative to develop a statistical framework to disentangle multiple introgression events involving more than two species from reciprocal introgression between two species, particularly because correctly identifying the species involved is essential for resolving introgression histories (Forsythe et al. 2025).

### Stochasticity in coalescent simulations

Our simulation results were largely consistent with our predictions for reciprocal introgression; however, we found that some windows that were simulated as reciprocally introgressed instead produced D-statistic results resembling non-introgressed tracts (see green points near the zero line in Fig. 2A). We observed this phenomenon to varying degrees in our replicate simulations (see Fig S1 for a run more heavily impacted). In most cases, the negative windows overcame the near-zero or positive values, however in 19/100 replicates, the average D-statistic across reciprocally introgressed windows was positive (Table S1), which is unexpected given our model (Fig 1). We believe that the unexpected near-zero reciprocal introgression windows result from the stepwise performance of the *msprime*. Our workflow involves simulating two separate gene flow events, P3->P2 followed by P2->P3. Although these events are set to occur at the same timing parameter, *msprime* treats them as stepwise events, meaning it’s possible that a subset of tracts transferred during the first introgression event could be immediately transferred back to the original donor during the second event, effectively undoing the transfer and rendering such an allele non-introgressed. This scenario differs from our definition of reciprocal introgression, which assumes that introgressed alleles mutually displace native alleles in both directions, yet our simulation workflow cannot distinguish between the two histories and lumps them together because both cases technically involve introgression in both directions. Given that this phenomenon is likely tied to an analytical quirk rather than a biological process and is negligible in most cases, we believe that our simulation strategy effectively models the genomic consequences of reciprocal introgression.

### Implications for empirical datasets

Our theoretical and simulation-based analyses highlight reciprocal introgression as an unaccounted-for outcome of gene flow, which has potential to bias and skew previous and future studies of introgression in empirical datasets. However, our study does not employ empirical genomic data, meaning an obvious open question that remains for future inquiry is whether reciprocal introgression can be detected in nature. We expect that the most likely scenarios for detecting reciprocal introgression will be in data that demonstrate bidirectional introgression. The best demonstrations of bidirectional introgression have been in systems that include population-level information (Yaakub et al. 2006; Wang et al. 2016; Pazmiño et al. 2019), meaning these may represent good starting points for exploring reciprocal introgression in empirical datasets.

Among the most compelling examples of hybridization are cases of adaptive introgression, in which beneficial alleles are obtained via gene flow (Dasmahapatra et al. 2012). While reciprocal introgression can occur via entirely neutral processes, our characterization of this novel form of introgression raises intriguing questions about whether there are scenarios in which natural selection would explicitly favor reciprocal introgression. One possible such scenario would be negative frequency-dependent selection (Ayala and Campbell 1974), in which newly introgressed alleles (initially infrequent in the population) are favored over the native alleles. If negative frequency-dependent selection acts in two populations for a given locus, then reciprocal displacement would be favored during the early generations following hybridization. As possible cases of reciprocal introgression are explored, it will be exciting to understand whether adaptation is involved. Together, our results move us toward a more complete understanding of the role of introgression in shaping genomes.

## Methods

Genome sequences were simulated using *msprime* (Baumdicker et al. 2022) and *tskit* (https://tskit.dev/) multispecies coalescent simulation packages in *Python* (v3.12.4). P3->P2 and P2->P3 introgression events were simulated separately at the same time point (40,000 generations before the present). Each introgression event was set to result in 20% of the genome being introgressed. The genomic coordinates of introgression tracts were retained as metadata and used to identify overlapping tracts (*i*.*e*., reciprocal introgression tracts). 100 replicate genome alignments were simulated, and representative examples are presented in Fig. 2 and Fig. S1. Full-genome alignments were split into 1,000 non-overlapping 10,000 bp windows, from which site patterns were counted and used to calculate D-statistics. See https://github.com/EvanForsythe/Reciprocal_introgression for the full set of parameters used to simulate genomes and for python scripts to reproduce all analyses.

## Supplemental material

**Table S1:** D-statistics from replicate simulations. Mean D-statistics for windows from introgression tract types are shown for 100 replicate simulation runs. “Expectations” indicates whether all D-statistic expectations (positive unidirectional introgression and negative reciprocal introgression) were met for a given run. ^*^ Run 81 was used for Fig. 2. ^*†*^ Run 24 was used for Fig. S1.

| Run | No int | P2 to P3 | P3 to P2 | Recip. Int. | Expectations |
| --- | --- | --- | --- | --- | --- |
| 1 | 0.032100742 | 0.690533045 | 0.839715795 | 0.20895845 | Failed |
| 2 | 0.007591654 | 0.39975405 | 0.842890373 | 0.10628515 | Failed |
| 3 | 0.043792117 | 0.592095622 | 0.857661497 | -0.0169468 | Passed |
| 4 | 0.054938687 | 0.837182262 | 0.874642085 | 0.18996004 | Failed |
| 5 | 0.037601465 | 0.687138112 | 0.884572288 | -0.505594 | Passed |
| 6 | 0.021691512 | 0.751871239 | 0.859907658 | -0.1395573 | Passed |
| 7 | 0.016672502 | 0.73783259 | 0.829213498 | -0.2381137 | Passed |
| 8 | 0.036622699 | 0.771635829 | 0.895965413 | -0.5440506 | Passed |
| 9 | 0.057567726 | 0.740160186 | 0.854414296 | -0.6251123 | Passed |
| 10 | 0.068160035 | 0.596047882 | 0.797722819 | -0.3221741 | Passed |
| 11 | 0.031968985 | -0.277327059 | 0.836532465 | -0.186932 | Failed |
| 12 | 0.011812271 | 0.457617315 | 0.838523191 | -0.3814852 | Passed |
| 13 | 0.006178703 | 0.724480394 | 0.794089347 | 0.24139826 | Failed |
| 14 | 0.106476837 | 0.721895817 | 0.826945744 | -0.539615 | Passed |
| 15 | 0.045048409 | 0.678253722 | 0.867619658 | -0.203349 | Passed |
| 16 | 0.021678736 | 0.616197137 | 0.861586434 | -0.6960011 | Passed |
| 17 | 0.002337404 | 0.683584607 | 0.851116046 | 0.41394833 | Failed |
| 18 | 0.023663906 | 0.661642589 | 0.698336082 | -0.228504 | Passed |
| 19 | 0.021817821 | 0.658319482 | 0.843187375 | -0.0367692 | Passed |
| 20 | 0.094132062 | 0.674388129 | 0.841361491 | -0.6233305 | Passed |
| 21 | 0.05257962 | 0.664715187 | 0.830168238 | -0.7341233 | Passed |
| 22 | 0.05257962 | 0.664715187 | 0.830168238 | -0.7341233 | Passed |
| 23 | 0.023582021 | 0.773286844 | 0.930483163 | -0.2915135 | Passed |
| 24<sup>†</sup> | 0.029711083 | 0.593978754 | 0.839811128 | 0.06049086 | Failed |
| 25 | 0.05072474 | 0.757320131 | 0.848434688 | -0.094035 | Passed |
| 26 | 0.030168003 | 0.705159192 | 0.831903268 | -0.7823687 | Passed |
| 27 | 0.028287322 | 0.553443873 | 0.890005625 | -0.0632551 | Passed |
| 28 | 0.081948808 | 0.758579699 | 0.771929761 | -0.0847095 | Passed |
| 29 | 0.033932731 | 0.420262325 | 0.838142361 | -0.0794322 | Passed |
| 30 | 0.011516493 | 0.591616079 | 0.767638937 | -0.3789553 | Passed |
| 31 | 0.031415134 | 0.663642845 | 0.883712024 | -0.8639695 | Passed |
| 32 | 0.012931843 | 0.720956571 | 0.876913403 | -0.7598743 | Passed |
| 33 | 0.019933347 | 0.714469321 | 0.856475867 | -0.7822377 | Passed |
| 34 | 0.046825421 | 0.66176317 | 0.824487928 | -0.3509978 | Passed |
| 35 | 0.040951125 | 0.595452258 | 0.872493253 | -0.5314738 | Passed |
| 36 | 0.025304727 | 0.85628671 | 0.882203732 | 0.03274585 | Failed |
| 37 | 0.036519958 | 0.698294582 | 0.880437644 | -0.0308203 | Passed |
| 38 | 0.061050172 | 0.681639502 | 0.881093565 | -0.155463 | Passed |
| 39 | 0.051981465 | 0.700041729 | 0.889224043 | 0.72353114 | Failed |
| 40 | 0.010193936 | 0.818044368 | 0.795384714 | -0.7492731 | Passed |
| 41 | 0.058294032 | 0.690936681 | 0.829195522 | -0.5583023 | Passed |
| 42 | 0.035019015 | 0.702354248 | 0.870285384 | 0.097215 | Failed |
| 43 | 0.027358663 | 0.597682765 | 0.824149829 | -0.6757483 | Passed |
| 44 | 0.081012768 | 0.648943319 | 0.751534515 | -0.6508205 | Passed |
| 45 | 0.049812027 | 0.760850904 | 0.86725288 | 0.26022717 | Failed |
| 46 | 0.028459971 | 0.835498811 | 0.854297818 | -0.4601488 | Passed |
| 47 | 0.0506803 | 0.649718051 | 0.859799842 | -0.5945335 | Passed |
| 48 | 0.004131084 | 0.8066228 | 0.782509448 | -0.2755792 | Passed |
| 49 | 0.025198944 | 0.746093851 | 0.87881936 | -0.5988393 | Passed |
| 50 | 0.020591778 | 0.712508679 | 0.91281775 | -0.3872705 | Passed |
| 51 | 0.017647246 | 0.657206564 | 0.588292666 | -0.5032426 | Passed |
| 52 | 0.039109648 | 0.734133553 | 0.840865424 | -0.8807159 | Passed |
| 53 | 0.000567012 | 0.834191233 | 0.687845849 | -0.8044942 | Passed |
| 54 | 0.031279155 | 0.742300813 | 0.881342227 | -0.6965079 | Passed |
| 55 | 0.062796689 | 0.689387653 | 0.827898227 | -0.2799494 | Passed |
| 56 | 0.018944184 | 0.563389076 | 0.884662015 | 0.36319031 | Failed |
| 57 | 0.046186396 | 0.768416131 | 0.858665898 | -0.7218563 | Passed |
| 58 | 0.021387648 | 0.733673512 | 0.906901856 | -0.3158441 | Passed |
| 59 | -0.003192595 | 0.735461739 | 0.895445606 | -0.3422183 | Passed |
| 60 | 0.040232534 | 0.56619515 | 0.787015635 | -0.257138 | Passed |
| 61 | 0.039271468 | 0.716012922 | 0.77088412 | -0.7420776 | Passed |
| 62 | 0.06284644 | 0.638763023 | 0.905809016 | 0.12683293 | Failed |
| 63 | 0.02165458 | 0.629437397 | 0.819373541 | -0.0253972 | Passed |
| 64 | 0.066566916 | 0.677779135 | 0.803975692 | -0.5158656 | Passed |
| 65 | 0.038610097 | 0.598252823 | 0.850379528 | -0.5998673 | Passed |
| 66 | 0.012233256 | 0.588904951 | 0.847033244 | -0.4957506 | Passed |
| 67 | 0.09003567 | 0.733266191 | 0.81911285 | -0.0443996 | Passed |
| 68 | 0.133750327 | 0.623841554 | 0.851774496 | -0.607169 | Passed |
| 69 | 0.046866943 | 0.797185247 | 0.874450502 | -0.5195052 | Passed |
| 70 | 0.041263787 | 0.672638601 | 0.916714053 | -0.366649 | Passed |
| 71 | 0.031252464 | 0.544070131 | 0.729473465 | 0.18107038 | Failed |
| 72 | 0.052866452 | 0.732938847 | 0.853843553 | -0.5749122 | Passed |
| 73 | 0.031999398 | 0.482896665 | 0.854179714 | -0.2084899 | Passed |
| 74 | 0.045092929 | 0.670241579 | 0.889854802 | 0.04213735 | Failed |
| 75 | 0.018092401 | 0.791651325 | 0.856366771 | -0.4579388 | Passed |
| 76 | 0.016415037 | 0.720296014 | 0.886767312 | -0.0339797 | Passed |
| 77 | 0.020904948 | 0.604573077 | 0.848663848 | -0.7124906 | Passed |
| 78 | 0.052854973 | 0.717700041 | 0.8027294 | -0.0540753 | Passed |
| 79 | 0.035723712 | 0.615925617 | 0.856353154 | -0.6525985 | Passed |
| 80 | 0.018320189 | 0.85397037 | 0.877558538 | 0.02482025 | Failed |
| 81* | 0.053689088 | 0.716182671 | 0.874483871 | -0.4552631 | Passed |
| 82 | 0.020802874 | 0.655766898 | 0.916160197 | 0.04305468 | Failed |
| 83 | 0.069211178 | 0.605222781 | 0.813060402 | -0.1637882 | Passed |
| 84 | 0.058915526 | 0.688355245 | 0.8648186 | -0.6693887 | Passed |
| 85 | 0.051598169 | 0.619829118 | 0.840540116 | -0.5212622 | Passed |
| 86 | 0.031544993 | 0.750364899 | 0.759941605 | -0.1025882 | Passed |
| 87 | 0.049945621 | 0.799517856 | 0.839501148 | -0.5120897 | Passed |
| 88 | 0.050744275 | 0.539392797 | 0.901428339 | 0.04946987 | Failed |
| 89 | 0.056488236 | 0.724042274 | 0.847091843 | -0.272629 | Passed |
| 90 | -0.000334852 | 0.807460676 | 0.870197421 | -0.3491208 | Passed |
| 91 | 0.029393733 | 0.574357876 | 0.832115298 | -0.1361068 | Passed |
| 92 | 0.053307864 | 0.817962873 | 0.909871746 | 0.03655685 | Failed |
| 93 | 0.043577064 | 0.702363595 | 0.873354358 | 0.0339347 | Failed |
| 94 | 0.052362557 | 0.760380976 | 0.789758516 | -0.0309391 | Passed |
| 95 | 0.030404933 | 0.689230582 | 0.827360751 | -0.5465571 | Passed |
| 96 | 0.038866065 | 0.465417438 | 0.86580692 | -0.7533442 | Passed |
| 97 | 0.055944395 | 0.690196148 | 0.806346555 | -0.7660179 | Passed |
| 98 | 0.044739508 | 0.821322962 | 0.885482305 | -0.0941885 | Passed |
| 99 | 0.074571627 | 0.487929209 | 0.889468136 | -0.0461959 | Passed |
| 100 | 0.022405541 | 0.741030755 | 0.863446324 | -0.2508651 | Passed |

**Figure S1:**
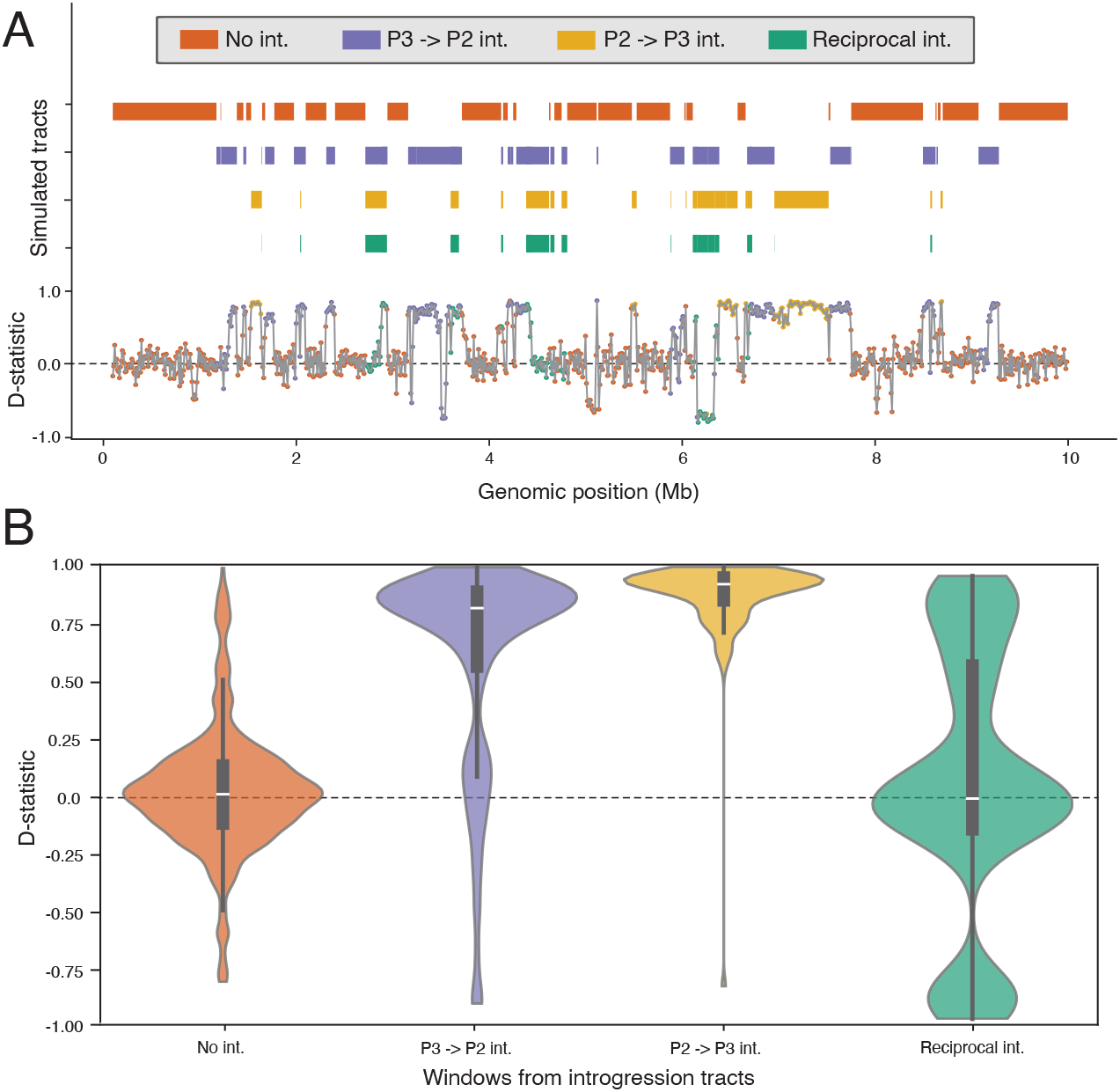
Example simulation run that did not fully adhere to reciprocal introgression expectations. (A) Simulated chromosome with tracts (i.e. haplotype blocks) that underwent alternative introgression histories. D-statistic estimates are shown for windows spanning the genome, which colored points indicated the introgression history of the window. (B) Violin plots showing the distributions of D-statistic estimates across windows of alternative introgression types. Box plots show medians and quartiles from distributions.

## Works cited

Ayala FJ and Campbell CA. Frequency-Dependent Selection.

Baumdicker F, Bisschop G, Goldstein D, Gower G, Ragsdale AP, Tsambos G, Zhu S, Eldon B, Ellerman EC, Galloway JG, et al. Efficient ancestry and mutation simulation with msprime 1.0. Genetics. 2022:220(3). 10.1093/genetics/iyab229

Crawford JE, Riehle MM, Guelbeogo WM, Gneme A, Sagnon N, Vernick KD, Nielsen R, and Lazzaro BP. Reticulate speciation and barriers to introgression in the anopheles gambiae species complex. Genome Biol Evol. 2015:7(11):3116–3131. 10.1093/gbe/evv203

Dagilis AJ, Peede D, Coughlan JM, Jofre GI, D’Agostino ERR, Mavengere H, Tate AD, and Matute DR. 15 years of introgression studies: quantifying gene flow across Eukaryotes 2021. 10.1101/2021.06.15.448399

Dasmahapatra KK, Walters JR, Briscoe AD, Davey JW, Whibley A, Nadeau NJ, Zimin AV, Hughes DST, Ferguson LC, Martin SH, et al. Butterfly genome reveals promiscuous exchange of mimicry adaptations among species. Nature. 2012:487(7405):94–98. 10.1038/nature11041

Durand EY, Patterson N, Reich D, and Slatkin M. Testing for Ancient Admixture between Closely Related Populations. Mol Biol Evol. 2011:28(8):2239–2252. 10.1093/molbev/msr048

Forsythe ES, Nelson ADL, and Beilstein MA. Biased gene retention in the face of introgression obscures species relationships. Genome Biol Evol. 2020a:12(9):1646–1663. 10.1093/GBE/EVAA149

Forsythe ES, Pappa BS, Clavette DA, and Mendoza DY. Detecting cryptic ghost lineage introgression in four-taxon genomic datasets. 2025. 10.1101/2025.04.28.651118

Forsythe ES, Sloan DB, and Beilstein MA. Divergence-based introgression polarization. Genome Biol Evol. 2020b:12(4):463–478. 10.1093/gbe/evaa053

Green RE, Krause J, Briggs AW, Maricic T, Stenzel U, Kircher M, Patterson N, Li H, Zhai W, Fritz MH-Y, et al. A Draft Sequence of the Neandertal Genome. Science (1979). 2010:328(5979):710–722. 10.1126/science.1188021

Hahn MW and Hibbins MS. A Three-Sample Test for Introgression. Mol Biol Evol. 2019:36(12):2878–2882. 10.1093/molbev/msz178

Hibbins MS and Hahn MW. The timing and direction of introgression under the multispecies network coalescent. Genetics. 2019:211(March):1059–1073.

Hibbins MS and Hahn MW. Phylogenomic approaches to detecting and characterizing introgression. Genetics. 2021. 10.1093/genetics/iyab173

Kuhlwilm M, Gronau I, Hubisz MJ, De Filippo C, Prado-Martinez J, Kircher M, Fu Q, Burbano HA, Lalueza-Fox C, De La Rasilla M, et al. Ancient gene flow from early modern humans into Eastern Neanderthals. Nature. 2016:530(7591):429–433. 10.1038/nature16544

Mallet J. Hybridization as an invasion of the genome. Trends Ecol Evol. 2005:20(5):229–237. 10.1016/j.tree.2005.02.010

Mallet J, Besansky N, and Hahn MW. How reticulated are species? BioEssays 2016:38(2):140–149. 10.1002/bies.201500149

Martin SH, Davey JW, and Jiggins CD. Evaluating the use of ABBA-BABA statistics to locate introgressed loci. Mol Biol Evol. 2015:32(1):244–257. 10.1093/molbev/msu269

Martin SH and Jiggins CD. Interpreting the genomic landscape of introgression. Curr Opin Genet Dev. 2017:47:69–74. 10.1016/j.gde.2017.08.007

Pazmiño DA, van Herderden L, Simpfendorfer CA, Junge C, Donnellan SC, Hoyos-Padilla EM, Duffy CAJ, Huveneers C, Gillanders BM, Butcher PA, et al. Introgressive hybridisation between two widespread sharks in the east Pacific region. Mol Phylogenet Evol. 2019:136:119–127. 10.1016/j.ympev.2019.04.013

Pease JB and Hahn MW. Detection and Polarization of Introgression in a Five-Taxon Phylogeny. Syst Biol. 2015:64(4):651–662. 10.1093/sysbio/syv023

Prüfer K, Racimo F, Patterson N, Jay F, Sankararaman S, Sawyer S, Heinze A, Renaud G, Sudmant PH, De Filippo C, et al. The complete genome sequence of a Neanderthal from the Altai Mountains. Nature. 2014:505(7481):43–49. 10.1038/nature12886

Rieseberg LH. Hybrid Speciation in Wild Sunflowers.

Rieseberg LH and Soltis DE. Phylogenetic consequences of cytoplasmic gene flow in plants. Evolutionary trends in Plants. 1991:5(1):65–84. 10.1007/s00606-006-0485-y

Suarez-Gonzalez A, Hefer CA, Christe C, Corea O, Lexer C, Cronk QCB, and Douglas CJ. Genomic and functional approaches reveal a case of adaptive introgression from Populus balsamifera (balsam poplar) in P.trichocarpa (black cottonwood). Mol Ecol. 2016:25(11):2427–2442. 10.1111/mec.13539

Taylor SA and Larson EL. Insights from genomes into the evolutionary importance and prevalence of hybridization in nature. Nat Ecol Evol. 2019:3(2):170–177. 10.1038/s41559-018-0777-y

Wang J, Street NR, Scofield DG, and Ingvarsson PK. Variation in Linked Selection and Recombination Drive Genomic Divergence during Allopatric Speciation of European and American Aspens. Mol Biol Evol. 2016:33(7):1754–1767. 10.1093/MOLBEV/MSW051

Yaakub SM, Bellwood DR, Herwerden L van, and Walsh FM. Hybridization in coral reef fishes: Introgression and bi-directional gene exchange in Thalassoma (family Labridae). Mol Phylogenet Evol. 2006:40(1):84–100. 10.1016/j.ympev.2006.02.012

Yakimowski SB and Rieseberg LH. The role of homoploid hybridization in evolution: A century of studies synthesizing genetics and ecology. Am J Bot. 2014:101(8):1247–1258. 10.3732/ajb.1400201

Zhang W, Dasmahapatra KK, Mallet J, Moreira GRP, and Kronforst MR. Genome-wide introgression among distantly related Heliconius butterfly species. Genome Biol. 2016:17(1). 10.1186/s13059-016-0889-0

Zheng Y and Janke A. Gene flow analysis method, the D-statistic, is robust in a wide parameter space. BMC Bioinformatics. 2018:19(1):1–19. 10.1186/s12859-017-2002-4

